# From Public Archive to Reusable Resource: Characterizing Gut Microbiome Metadata in the NCBI SRA

**DOI:** 10.64898/2026.08.31.742652

**Authors:** Kiersten A. Oderman, Tarak N. Nandi, Judy Malas, Ravi K. Madduri, Jarrad T. Hampton-Marcell

## Abstract

Public sequencing repositories contain large amounts of gut microbiome data that could support cross-study comparison, reproducibility analysis, and microbiome foundation model development. However, the extent to which these data are structured, harmonized, and reusable at archive scale remains unclear. Here, we characterized publicly available gut microbiome sequencing metadata from the NCBI Sequence Read Archive using Google BigQuery, focusing on human gut metagenome, mouse gut metagenome, and broadly annotated gut metagenome records. We evaluated temporal growth, sequencing depth, BioSample and BioProject structure, platform and instrument use, metadata completeness, host attribution, publication linkage, and research themes from linked literature. Public gut microbiome data increased substantially over time and were dominated by human-associated datasets and Illumina sequencing platforms. Core technical metadata fields were highly complete, but biological context needed for reuse, including host identity, phenotype, study design, and disease status, was often inconsistently encoded or required recovery from BioSample attributes and linked publications. In the generic “gut metagenome” cohort, host identity could be assigned for only 13.00% of BioSamples, highlighting the limitations of broad organism annotations for automated cohort construction. Publication linkage was also incomplete at the archive level, although usable text was recovered for most linked publications. Topic modeling of SRA-linked literature showed persistent emphasis on core gut microbiota composition and increasing representation of human cohort and infant microbiome studies. Overall, these findings show that public gut microbiome data are extensive and technically rich but not uniformly analysis ready. Improved metadata harmonization, publication linkage, and biological context recovery will be necessary to support reliable large-scale reuse and AI-ready microbiome data resources.

## Introduction

As biological data generation has increased, public repositories such as the National Center for Biotechnology Information (NCBI) Sequence Read Archive (SRA) have become essential for storing and sharing sequencing datasets. Containing tens of petabytes of raw sequence data from human, microbial, animal, and environmental studies, the SRA provides the scale needed for data reuse, cross-study comparison, and large-scale computational analysis. The broader NCBI ecosystem includes more than 40 interconnected databases spanning genomic sequences, scientific literature, protein structures, and clinical variants, making it a central infrastructure for modern biological research. Systematic reanalysis of archived data can reveal patterns across studies, test the reproducibility of published findings, and support questions that cannot be addressed within individual experiments. In gut microbiome research, for example, public sequencing datasets have been used to compare microbial patterns across populations, host systems, and disease states, including cross-disease meta-analyses, curated metagenomic resources, and large population-scale microbiome cohorts^1–3^. NCBI is particularly valuable for this type of work because of its large scale, standardized sequence formats, integrated tools such as BLAST and the SRA Toolkit, and continued curation.

Gut microbiome research connects microbial ecology with many areas of human health and disease. The gut microbiome is no longer studied only as a feature of gastrointestinal biology; it is now relevant to metabolism, immune regulation, cancer, cardiovascular disease, nutrition, pharmacology, and host-microbe interactions more broadly. This broad relevance has made gut microbiome data valuable across multiple disciplines, including microbiology, immunology, metabolomics, medicine, epidemiology, bioinformatics, and machine learning methods are being used to identify microbial biomarkers, predict disease risk, and design targeted microbiome-based therapies. As sequencing-based microbiome studies continue to accumulate, public repositories now contain a large and diverse body of gut microbiome data that could support cross-study comparisons, reproducibility analyses, and large-scale computational modeling. However, the proportion of the NCBI SRA that has been successfully reused or reanalyzed has not been comprehensively quantified. Prior work has nevertheless identified several barriers to secondary use, including incomplete or inconsistent metadata, uneven annotation practices, difficulty determining whether datasets are suitable for a new analysis, and the technical demands of processing large-scale sequencing archives^4–6^. As a result, the field faces a central challenge: not a lack of microbiome data, but a lack of sufficiently standardized and interoperable information needed to identify and integrate relevant datasets.

These barriers represent fundamental challenges at the intersection of biology and data science. Community standards have been developed to improve how sequencing studies are described, including the Minimum Information about any (x) Sequence (MIxS) specifications for reporting sample and sequence metadata, and the Strengthening the Organization and Reporting of Microbiome Studies (STORMS) checklist for human microbiome research^7,8^. More recently, the Standards for Technical Reporting in Environmental and Host-Associated Microbiome Studies (STREAMS) guidelines expanded this framework to environmental, synthetic, and non-human host-associated microbiome studies^9^. However, these standards are not applied consistently across all submissions, and repositories may still accept metadata entered through free-text fields or project-specific terminology. As a result, functionally identical information may be recorded using different terminologies, units, or ontological frameworks across datasets, making automated filtering and comparison difficult without extensive manual curation^5,10^. For example, stool-derived gut microbiome samples may be described as “stool,” “feces,” “fecal,” “faecal,” “gut,” “intestinal contents,” or simply “human gut metagenome,” even when they refer to biologically similar sample types. Without harmonization, these inconsistencies can fragment otherwise comparable datasets and complicate cohort construction for downstream analyses.

Beyond metadata challenges, the computational requirements for large-scale reanalysis are substantial. Researchers need proficiency in command-line interfaces, high-performance computing environments, bioinformatics pipelines, and programming languages to download, process, and analyze millions of sequencing reads. Meaningful interpretation also requires expertise in statistics, machine learning, and domain-specific biology. In practice, many researchers are trained deeply in their biological discipline but have fewer opportunities for formal training in bioinformatics, programming, and large-scale data analysis, making public sequencing data difficult to reuse without additional computational support^11^. This skills gap creates a critical bottleneck: while biologists possess the domain expertise to formulate important research questions and interpret biological significance, they may lack consistent access to the computational tools, infrastructure, or workflows needed to analyze the datasets that could answer those questions. Consequently, publicly funded sequencing data remain underused, limiting opportunities to validate findings, generate new hypotheses, and support translational discovery.

This divide between biological expertise and computational proficiency highlights the need for computational frameworks that can interpret biological context while operating across complex and heterogeneous data environments. AI foundation models offer one promising approach. These large-scale models learn generalizable representations from diverse datasets and can then be adapted to multiple downstream tasks, making them especially useful in data scarce settings. In biological research, they may help connect information across fragmented metadata structures, experimental platforms, and databases. Robust foundation models could also support agentic AI systems capable of searching, harmonizing, interpreting, and reasoning across biological data resources. Agentic AI refers to autonomous software systems that can perceive, reason, plan, and act with limited human intervention to complete complex, goal-oriented tasks^12^. Although these systems have shown promise in other domains requiring large-scale data integration and pattern recognition, their application to microbiome research remains limited by the lack of domain-specific foundation models for host-microbe interactions^13^. Unlike general-purpose language models trained primarily on text, microbiome foundation models must represent relationships among microbial communities, host phenotypes, environmental exposures, and disease-associated variation. This requires a clear understanding of the structure, consistency, and interoperability of the available microbiome data.

The usefulness of future AI systems in gut microbiome research will therefore depend heavily on the quality and organization of the data used to train them. Foundation models are not a substitute for curation; rather, they require structured and biologically meaningful input data to learn reliable representations. Before specialized foundation models for host-microbe interactions can be developed, the current landscape of publicly available gut microbiome datasets must be characterized. This includes determining what data are available, how they are organized, how complete their metadata are, and which biological and methodological areas are represented. In this study, we systematically characterize gut microbiome datasets in the NCBI SRA through large-scale metadata analysis and topic modeling of linked publications. By defining the structure, scope, and current limitations of the archive, this work establishes a starting point for microbiome data integration and future foundation model development.

## Methods

### Data Retrieval and Cohort Definition

Publicly available gut microbiome sequencing metadata were retrieved from the National Center for Biotechnology Information Sequence Read Archive (NCBI SRA) using the NCBI SRA metadata table hosted in Google Cloud BigQuery. BigQuery was selected because the analysis required structured filtering and aggregation across more than one million sequencing records. The SRA BigQuery resource supports archive-scale querying and parallel data extraction, whereas NCBI Entrez is more appropriate for targeted searches and record-level retrieval and is subject to request-rate limitations. Entrez was therefore used for linking SRA records to associated publications, while BigQuery was used for the primary metadata analysis.

Records were identified using keyword-based queries of the SRA organism field for gut-associated annotations containing terms related to microbiome, metagenome, stool, or feces (**Figure 1**). Retrieved records were then restricted to the three most highly represented organism annotations: “human gut metagenome,” “mouse gut metagenome,” and “gut metagenome.” These records were analyzed as the human-associated, mouse-associated, and generic cohorts, respectively. Records annotated as “gut metagenome” were retained as a separate generic cohort and were later examined using BioSample metadata to infer host identity. All analyses were restricted to publicly available sequencing runs.

**Figure 1.**
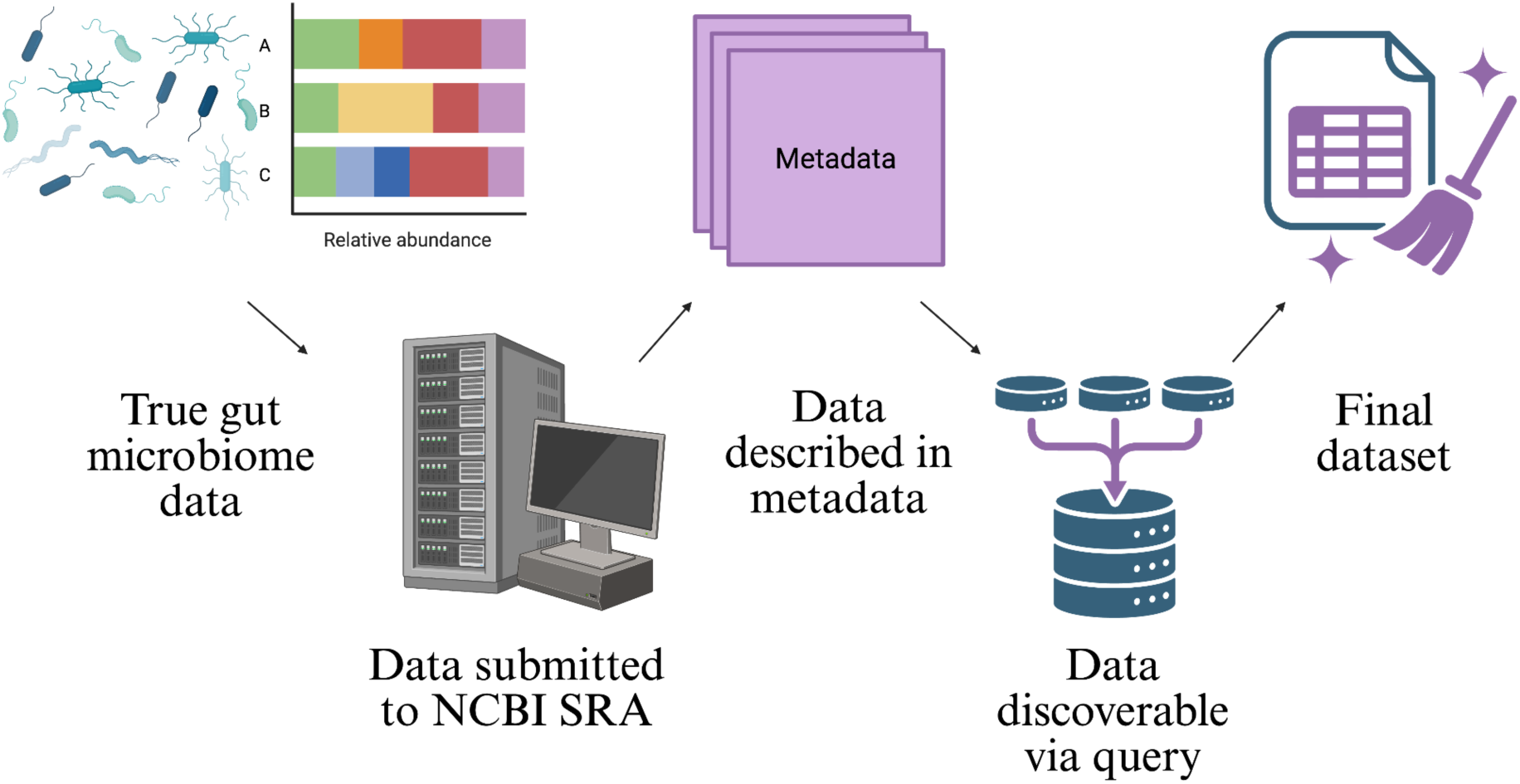
Overview of Data Retrieval Procedures.

### Metadata Processing and Dataset Structure

Metadata were extracted at the sequencing-run level and included accession identifiers, release date, organism annotation, BioProject accession, BioSample accession, SRA Study accession, sequencing depth, library preparation descriptors, platform, instrument, and assay type. Release dates were converted to calendar year, and character fields were standardized to reduce differences in capitalization, punctuation, and spacing.

The SRA is organized hierarchically, with individual sequencing runs representing technical sequencing files generated from biological samples. Multiple sequencing runs can be associated with the same BioSample, and multiple BioSamples can be grouped within a BioProject, which represents the larger study or research project. Using these accession relationships, the hierarchical structure of the SRA was reconstructed by linking sequencing runs to their associated BioSamples and BioProjects. Runs per BioSample and runs per BioProject were calculated separately for the human, mouse, and generic cohorts. Distributions were summarized using the median, interquartile range, maximum, percentage of units represented by a single run, and the proportion of total runs contributed by the top 1% of BioSamples or BioProjects.

### Temporal Growth Analysis

Sequencing runs were aggregated by year of release to evaluate changes in data availability over time. Annual counts were calculated separately for the human, mouse, and generic cohorts, and records released in 2026 were excluded because the year was incomplete at the time of analysis. Temporal trends were assessed using Spearman rank correlation, with calendar year treated as the independent ordered variable and the annual metric of interest treated as the dependent variable. The Spearman correlation coefficient (ρ) was used to describe the direction and strength of monotonic change over time, and two-sided p-values were used to assess statistical significance at α = 0.05.

### Sequencing Depth Analysis

Sequencing depth was evaluated using the total megabases reported per sequencing run. Records with missing or nonpositive sequencing-depth values were excluded from this analysis. For each cohort and year, the median sequencing depth and interquartile range were calculated to represent central tendency and variability in sequencing effort.

Temporal changes in sequencing depth were evaluated using Spearman rank correlations between year and annual median sequencing depth. Changes in variability were similarly evaluated using correlations between year and annual IQR width. To provide an interpretable comparison between earlier and more recent studies, mean annual median depth and mean annual IQR width were also compared between records released in 2014 or earlier and those released from 2021 through 2025.

### Platform and Instrument Analysis

Sequencing platform names and instrument models were standardized before analysis. Platform use was summarized as the annual proportion of runs generated using the most frequently represented technologies, including Illumina, LS454, Ion Torrent, PacBio SMRT, and Oxford Nanopore. Overall platform proportions, first year of appearance, peak year, and peak annual proportion were calculated. Spearman rank correlations were used to evaluate monotonic changes in annual platform representation through 2025.

Instrument models were grouped into broader instrument families, including 454/GS FLX, MiSeq, HiSeq, NovaSeq, NextSeq, Ion Torrent, and iSeq. Instrument use was calculated as the annual proportion of runs represented by each instrument family. Trend analyses were performed using Spearman rank correlations between year and annual instrument proportion.

Instrument patterns were also summarized across three periods, 2010–2015, 2015–2020, and 2020–2025, to characterize the transition from early lower-throughput systems to newer high-throughput instruments.

### Metadata Completeness, Consistency, Biological Richness, and Interoperability

Metadata completeness was evaluated for the run-level fields reported in **Supplemental Figure 1**. These fields included experiment accession, organism name, instrument, study accession, sample accession, total size in megabases, library name, library strategy, library source, and library selection. For each field, values were classified as missing if they were stored as null values, empty strings, or common placeholder terms, including “unknown,” “not available,” “not provided,” “none,” or related missing-value labels. Missingness was calculated separately for the human, mouse, and generic gut metagenome cohorts as the number of missing records divided by the total number of records in that cohort and multiplied by 100. Field definitions and cohort-specific missingness percentages are reported in **Supplemental Figure 1**.

Because field completeness does not necessarily indicate that metadata are standardized or biologically informative, additional analyses were performed to assess reporting consistency, biological richness, and interoperability. Reporting consistency was evaluated by examining populated technical metadata fields before and after formatting normalization. Values were standardized by normalizing capitalization, punctuation, whitespace, and separators. For each field and cohort, raw unique values and normalized unique values were compared to estimate how much apparent value diversity was due to formatting differences.

Biological richness was evaluated using host attribution in the generic “gut metagenome” cohort. Available BioSample metadata were parsed using rule-based classification of scientific names, taxonomic identifiers, and free-text attributes. Records were classified as assigned when a host category could be inferred, unknown when BioSample metadata were available but insufficient for host assignment, and unmapped when BioSample attributes could not be retrieved. Host-attribution success was summarized as both the overall percentage of generic BioSamples assigned to a host and the percentage assigned among records with retrieved BioSample metadata.

Interoperability was evaluated by measuring linkage between SRA records and associated publications. BioProject and SRA Study accessions were mapped to PubMed identifiers using NCBI Entrez through BioPython. PubMed identifiers were then mapped to PubMed Central identifiers when available, and publication text availability was summarized as full text, abstract only, or no usable text. These publication-linkage and text-recovery metrics were used to assess the extent to which biological context could be recovered beyond the structured run-level metadata.

### Topic Modeling of Linked Literature

Publications associated with the selected SRA records were identified using BioProject and SRA Study accessions and retrieved through the NCBI Entrez API using BioPython. Duplicate PubMed identifiers were removed, resulting in 718 unique linked publications. PubMed identifiers were mapped to PubMed Central identifiers when available, and full text was used when accessible; abstracts were used when full text was unavailable. The final hybrid corpus included 717 documents, consisting of 526 full-text articles and 191 abstract-only records.

Text was converted to lowercase, stripped of punctuation, and tokenized. General English stop words and corpus-specific terms that primarily reflected article structure rather than biological content, such as “figure,” “table,” and calendar years, were removed. Documents with insufficient text after preprocessing were excluded. Topic modeling was performed using the tomotopy Python package with a 20-topic Dirichlet Multinomial Regression model and publication year included as document-level metadata.

A 20-topic model was selected to provide sufficient thematic resolution while retaining interpretability across the linked literature corpus. Final topic labels were assigned through manual review of the top-ranked words for each topic and comparison with representative documents showing high topic proportions. Topics were retained when they represented distinct and interpretable biological or methodological themes. Topics dominated by formatting terms, publication boilerplate, or overlapping vocabulary were refined through additional stop-word filtering and model refitting. This decision process was used to balance statistical separation with biological interpretability.

### Topic-Platform and Topic-Host Associations

Topic prevalence estimates were linked to annual platform-use proportions to examine relationships between research themes and sequencing technologies. Topic-platform association values were calculated by combining annual topic prevalence with the annual proportional representation of each sequencing platform. These values represent period-level associations between topic prevalence and platform use, rather than direct causal relationships between individual publications and sequencing technologies. Associations were visualized using heatmaps and Sankey diagrams.

For visualization and temporal comparison, topic prevalence and topic-platform analyses were restricted to publications from 2008 through 2025 to match the primary SRA metadata analysis window and reduce the influence of sparse earlier publication-year values. To evaluate changes over time, topic-platform associations were summarized across four non-overlapping periods: 2005-2009, 2010-2014, 2015-2019, and 2020-2025. Topic distributions were also compared across human, mouse, and generic cohorts to evaluate relationships between biological study systems and research focus.

### Statistical Analysis

Runs per BioSample and BioProject were strongly right-skewed; therefore, differences among the human, mouse, and generic cohorts were evaluated using Kruskal-Wallis tests. Spearman rank correlations were used to assess temporal trends in sequencing depth, sequencing-depth variability, platform use, instrument use, and topic prevalence. Statistical significance was evaluated using an alpha level of 0.05. Because the primary objective was descriptive characterization of the SRA archive, effect sizes, proportions, medians, interquartile ranges, and temporal trends were emphasized alongside hypothesis-test results.

### Computational Environment

All analyses were performed in Python 3.12.13 on a Linux-based Google Colab environment. Data processing, statistical analysis, topic modeling, and visualization were conducted using pandas v2.2.2, NumPy v2.0.2, SciPy v1.16.3, Matplotlib v3.10.0, Plotly v5.24.1, tomotopy v0.14.0, BioPython v1.87, and google-cloud-bigquery v3.42.1^14–19^. The primary metadata analysis used the NCBI SRA metadata table hosted in Google Cloud BigQuery. Publication linkage was performed using the NCBI Entrez API through BioPython, with PubMed and PubMed Central records retrieved for linked literature analysis^20^. Because the SRA, PubMed, and PubMed Central are continuously updated resources, all analyses represent a snapshot of publicly available records at the time of retrieval. Figures were exported at 600 dpi and saved to a dedicated Google Drive output directory. Analysis code and notebooks used to generate figures and summary statistics are available at https://github.com/kierstenoderman/SRA_characterization.

### Data and Code Availability

All analyses used publicly available metadata from the NCBI Sequence Read Archive, BioSample, BioProject, PubMed, and PubMed Central. Analysis code used for metadata processing, statistical analysis, topic modeling, visualization, and generation of summary tables is available at: https://github.com/kierstenoderman/SRA_characterization.

The repository includes the final analysis notebook used to generate publication-ready figures and Results summary statistics. Large intermediate metadata files are not stored in the repository because of file size constraints, but the notebook describes the expected input directory structure and public data sources needed to reproduce the analysis.

## Results

### Overview of Retrieved Gut Microbiome Sequencing Metadata

A search for “gut microbiome” in the NCBI Sequence Read Archive returned over 1.1 million public sequencing records, spanning both metagenomic and amplicon-based studies. Publicly archived gut microbiome sequencing data were dominated by short-read platforms, with more than 1.12 million reads generated on Illumina instruments and an additional 5,386 generated on BGISeq platforms. Long-read technologies were comparatively underrepresented, including 11,848 Oxford Nanopore and 11,227 PacBio SMRT records. Major organism labels included gut metagenome (599,636), human gut metagenome (274,882), and mouse gut metagenome (69,011), reflecting both the breadth of host systems represented and the persistence of broad organism annotations within public repositories.

A dataset of publicly available gut microbiome sequencing metadata was retrieved from the NCBI Sequence Read Archive (SRA) using targeted keyword queries. Records were initially examined by organism annotation to characterize the distribution of host-associated datasets. Across all retrieved records, human-associated samples dominated the dataset, comprising over 1.1 million sequencing runs and accounting for over 45% of the total data, followed by generic “gut metagenome” entries (∼579,000 runs) and mouse-associated samples (∼340,000 runs). Additional host categories, including livestock (e.g., pig and bovine), poultry, aquatic organisms, and other animal models, were represented at substantially lower frequencies (**Supplemental Figure 2**). Because public repositories such as the SRA are continuously updated as new studies are deposited, the structure and composition of available gut microbiome data are not static. This ongoing growth highlights the need for scalable computational and AI-assisted approaches that can repeatedly identify, harmonize, and interpret newly available datasets.

This distribution highlights a strong bias toward human microbiome research within publicly archived sequencing data, with comparatively fewer datasets derived from model organisms and non-mammalian hosts. The presence of a large “generic” gut metagenome category further indicates inconsistencies in host annotation, which may obscure biological interpretation and complicate downstream analyses. Initial inspection also confirmed that SRA metadata are structured around sequencing runs rather than biological samples or studies, necessitating hierarchical analysis across runs, BioSamples, and BioProjects to accurately characterize dataset organization and evaluate data reuse potential. For downstream analysis, we focused on the largest organism annotations, human gut metagenome (45.66%), gut metagenome (23.51%), and mouse gut metagenome (13.85%) which together accounted for 83.02% of retrieved records. These groups were selected because they captured the majority of available gut microbiome data while retaining biological meaningful comparisons between human-associated and broadly annotated gut metagenome records.

### Growth of Publicly Available Gut Microbiome Sequencing Data

The number of publicly archived gut microbiome sequencing runs increased markedly over time across all organism groups, with particularly rapid expansion observed after approximately 2014 (**Figure 2a**). Human-associated datasets showed the most pronounced growth, rising sharply through the late 2010s. Generic “gut metagenome” datasets followed a similar trajectory but at lower overall magnitude, while mouse-associated datasets demonstrated steady but more moderate growth throughout the same period.

**Figure 2.**
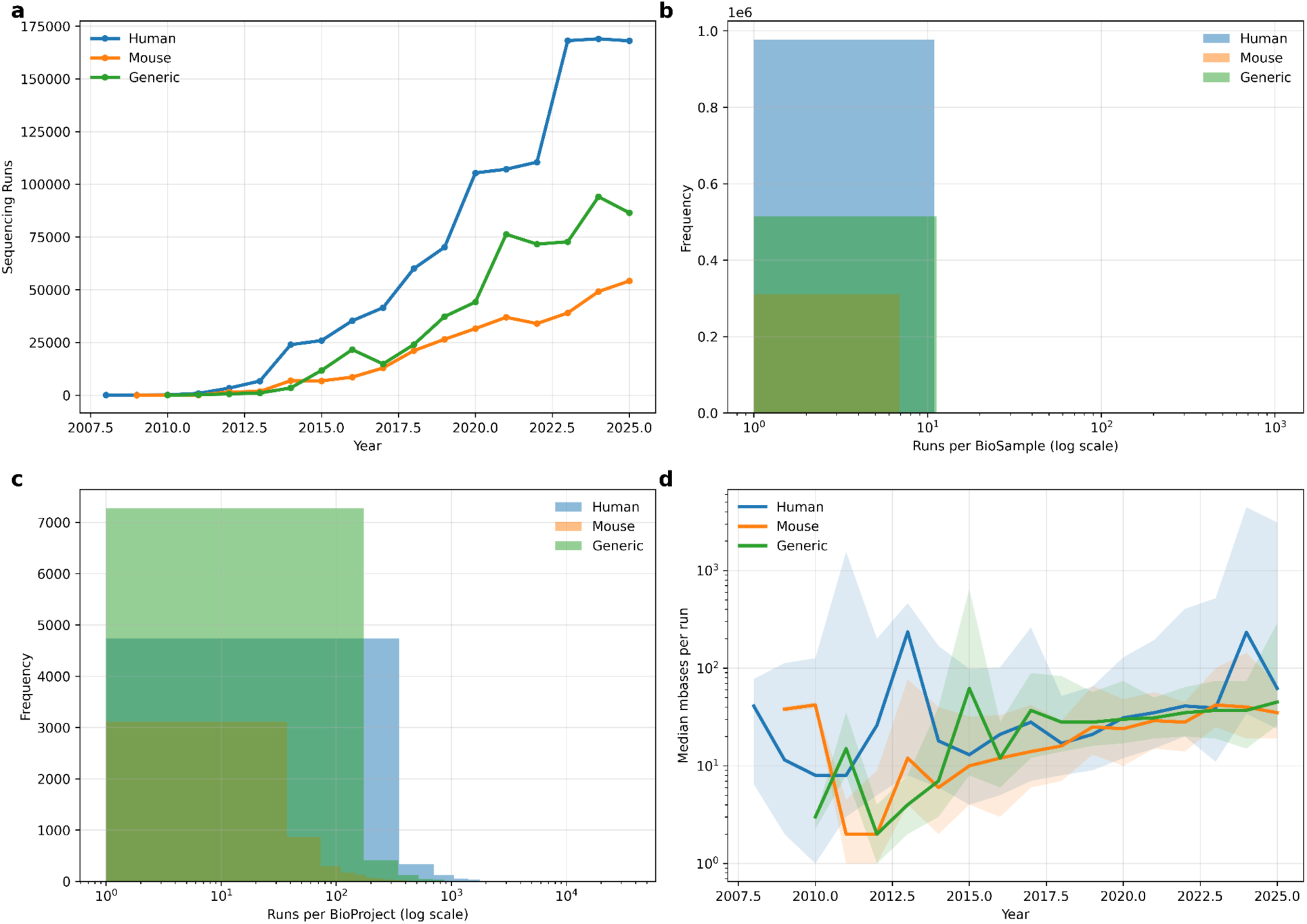
Temporal growth, dataset structure, and sequencing depth of public gut microbiome SRA records. (**a**) Annual number of human, mouse, and generic gut microbiome sequencing runs through 2025. (**b**) Distribution of sequencing runs per BioSample. (**c**) Distribution of sequencing runs per BioProject. (**d**) Median sequencing depth per run with interquartile ranges across cohorts through 2025. Log-scaled axes were used in panels b–d to visualize strongly skewed distributions and large differences in sequencing depth.

Notably, all cohorts exhibited sustained increases over time, reflecting the widespread adoption of high-throughput sequencing technologies and growing interest in microbiome research. Analyses excluded incomplete 2026 data to avoid artifacts from partial reporting, and trends remained consistent when restricted to fully reported years. Collectively, these findings demonstrate a sustained and accelerating expansion of publicly available gut microbiome sequencing resources over the past decade.

### Hierarchical Structure of SRA Data Representation

Across all organism groups, the majority of BioSamples were represented by a single sequencing run (median = 1; **Figure 2b**). More than 93% of BioSamples in each cohort contained only one run. However, the distributions were strongly right-skewed, with maximum values ranging from 592 runs in the mouse cohort to 1,030 runs in the generic cohort. This pattern indicates that most BioSamples were sequenced once, while a small subset was associated with extensive replication or repeated measurements.

Project-level analysis revealed substantial concentration of sequencing effort (**Figure 2c**). Human-associated BioProjects contained a median of 73 runs per project, compared with 28 runs for mouse projects and 27 runs for generic projects. The top 1% of projects contributed approximately 20–28% of all runs across cohorts, with the largest concentration observed among human-associated projects. These findings show that a relatively small number of large studies contribute disproportionately to the public archive.

Median sequencing depth increased over time across all organism groups, with the largest changes occurring in more recent years (**Figure 2d**). Comparing records from 2014 or earlier with those from 2021–2025, mean annual median sequencing depth increased approximately 1.7-fold in human datasets, 2.0-fold in mouse datasets, and 6.0-fold in generic gut metagenome datasets. Sequencing depth was positively associated with year in the human and generic cohorts, while the mouse cohort showed a similar but weaker trend.

Mouse and generic gut metagenome datasets showed more gradual increases in sequencing depth, whereas human-associated datasets displayed greater year-to-year variability. Interquartile ranges also widened over time, particularly in mouse and generic datasets, indicating increasing heterogeneity in sequencing depth and experimental design. Together, these patterns suggest a shift toward deeper and more varied sequencing strategies as gut microbiome studies expanded in scale and purpose.

### Platform and Instrument Trends in Gut Microbiome Research

Analysis of sequencing platforms revealed a strong and increasing dominance of Illumina technologies across all cohorts (Figure 3). Overall, 91.82% of sequencing runs were generated on Illumina platforms, with the annual proportion reaching 97.04% in 2024. Illumina adoption increased strongly over time (Spearman ρ = 0.89, p < 0.001), while LS454 showed the opposite pattern, declining sharply from its early predominance (ρ = −1.00, p < 0.001). Smaller contributions from Ion Torrent, PacBio, and Oxford Nanopore were also observed. Although PacBio and Oxford Nanopore each accounted for less than 1% of all runs overall, both increased significantly over time and reached their highest annual representation in 2025. Together, these trends show a clear transition from early LS454-based sequencing to broad Illumina dominance, with gradual diversification toward newer long-read technologies.

**Figure 3.**
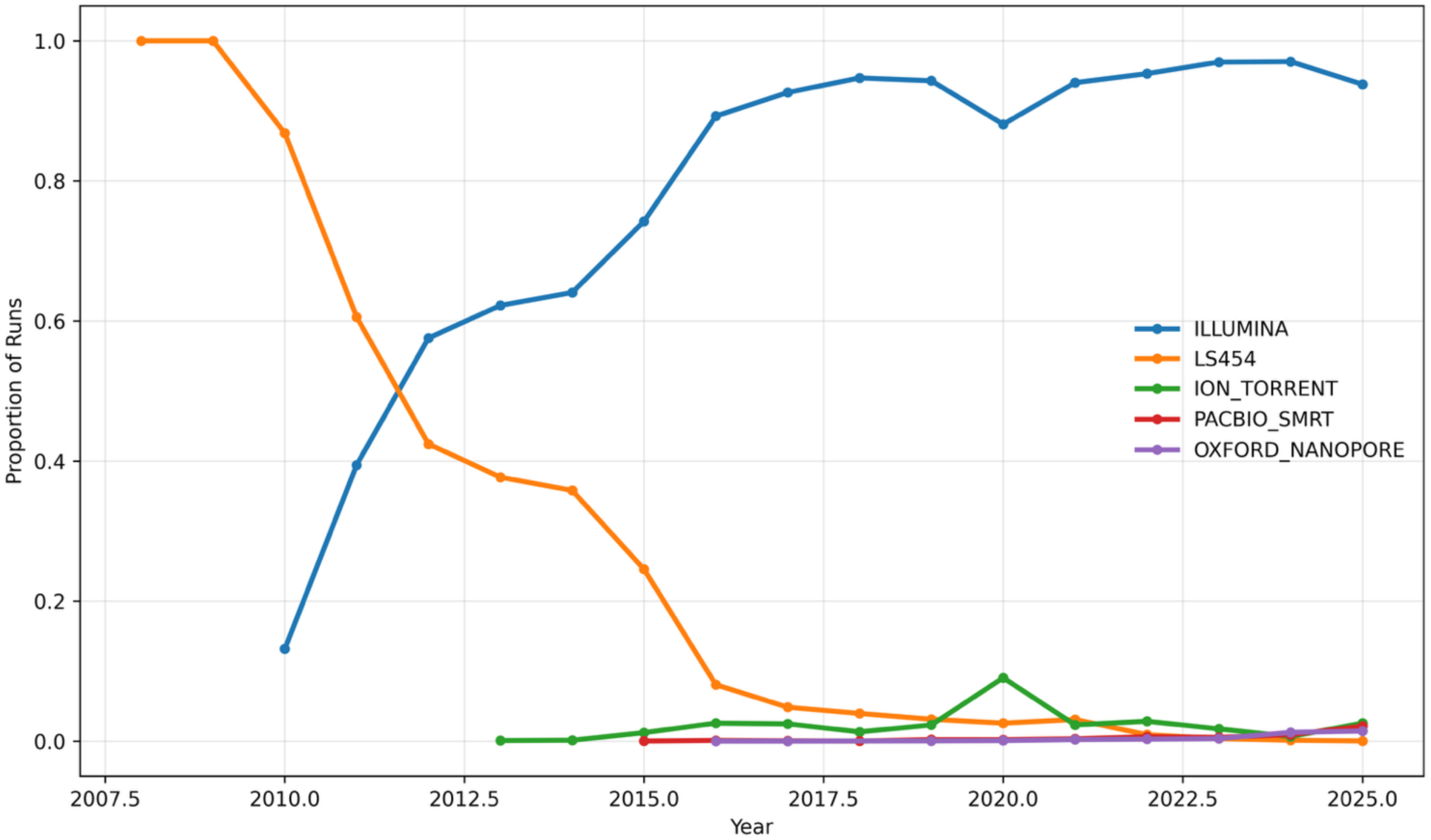
Sequencing Platform Adoption Over Time (All Cohorts, Through 2025).

Instrument-level analysis showed a clear transition from early 454-based sequencing to successive generations of Illumina instruments (**Figure 4**). The 454/GS FLX platform accounted for nearly all runs among the major instruments from 2008 through 2011, but its use declined sharply thereafter (Spearman ρ = −0.99, p < 0.001). HiSeq became prominent during the early transition period, reaching its highest annual proportion in 2013 at 39.58% of runs. MiSeq subsequently became the most widely used instrument, accounting for 53.26% of runs overall and reaching a peak annual proportion of 70.63% in 2018. Its use remained substantial through 2025, although the overall temporal trend was not monotonic because its proportional share declined as newer instruments entered the archive.

**Figure 4.**
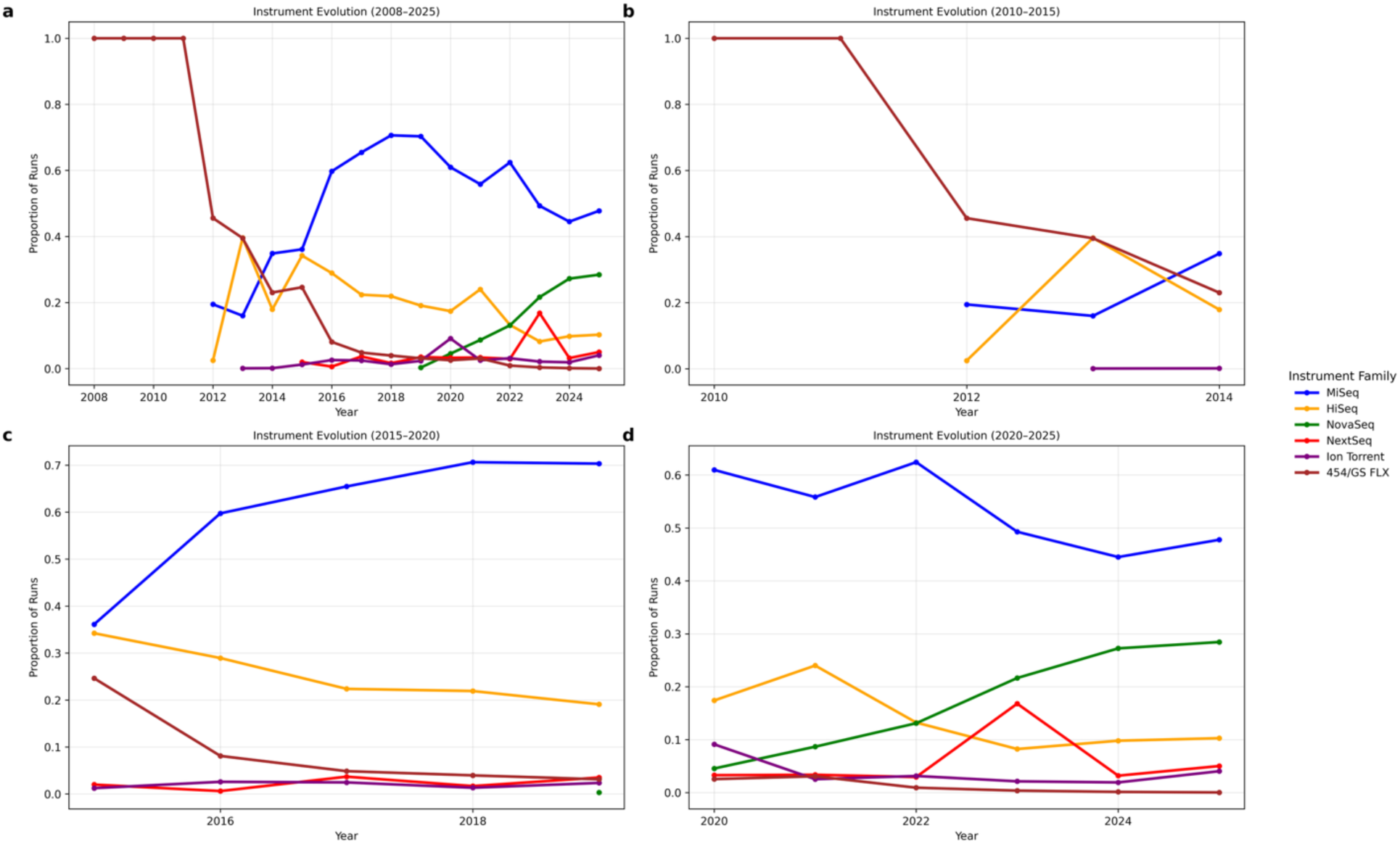
Temporal evolution of major sequencing instruments in public gut microbiome datasets. (**a**) Overall proportional trends in major sequencing instruments across all cohorts through 2025. (**b**) Instrument trends during the early transition period (2010–2015). (**c**) Instrument trends during the intermediate adoption period (2015–2020). (**d**) Instrument trends during the recent high-throughput period (2020–2025).

The most recent transition was marked by the emergence of NovaSeq. NovaSeq first appeared in 2019, exceeded 10% of runs among the major instruments by 2022, and increased to 28.44% in 2025. This increase was strongly monotonic over time (Spearman ρ = 1.00, p < 0.001). Together, these trends identify three broad periods of instrument adoption: early reliance on 454/GS FLX, expansion of HiSeq and MiSeq between 2012 and 2020, and increasing use of NovaSeq after 2020. These shifts reflect the movement toward higher-throughput sequencing systems capable of supporting larger cohorts and deeper microbiome profiling.

### Metadata Completeness and Reporting Consistency

Metadata completeness was evaluated across the run-level fields reported in **Supplemental Figure 1**. Overall, core technical metadata were highly complete across the human, mouse, and generic gut metagenome cohorts (**Figure 5a**). Experiment accession, organism name, instrument, sample accession, library strategy, library source, and library selection were present for all records in the analyzed dataset. Study accession showed very low missingness across cohorts, ranging from 0.02% to 0.03%. Total sequencing size showed slightly higher missingness, with 0.48% missingness in human records, 0.87% in mouse records, and 0.48% in generic records. Library name was the least complete field, with missingness ranging from 3.84% in generic records to 11.79% in human-associated records. These results indicate that the SRA captures the basic technical information needed to identify and process sequencing runs.

**Figure 5.**
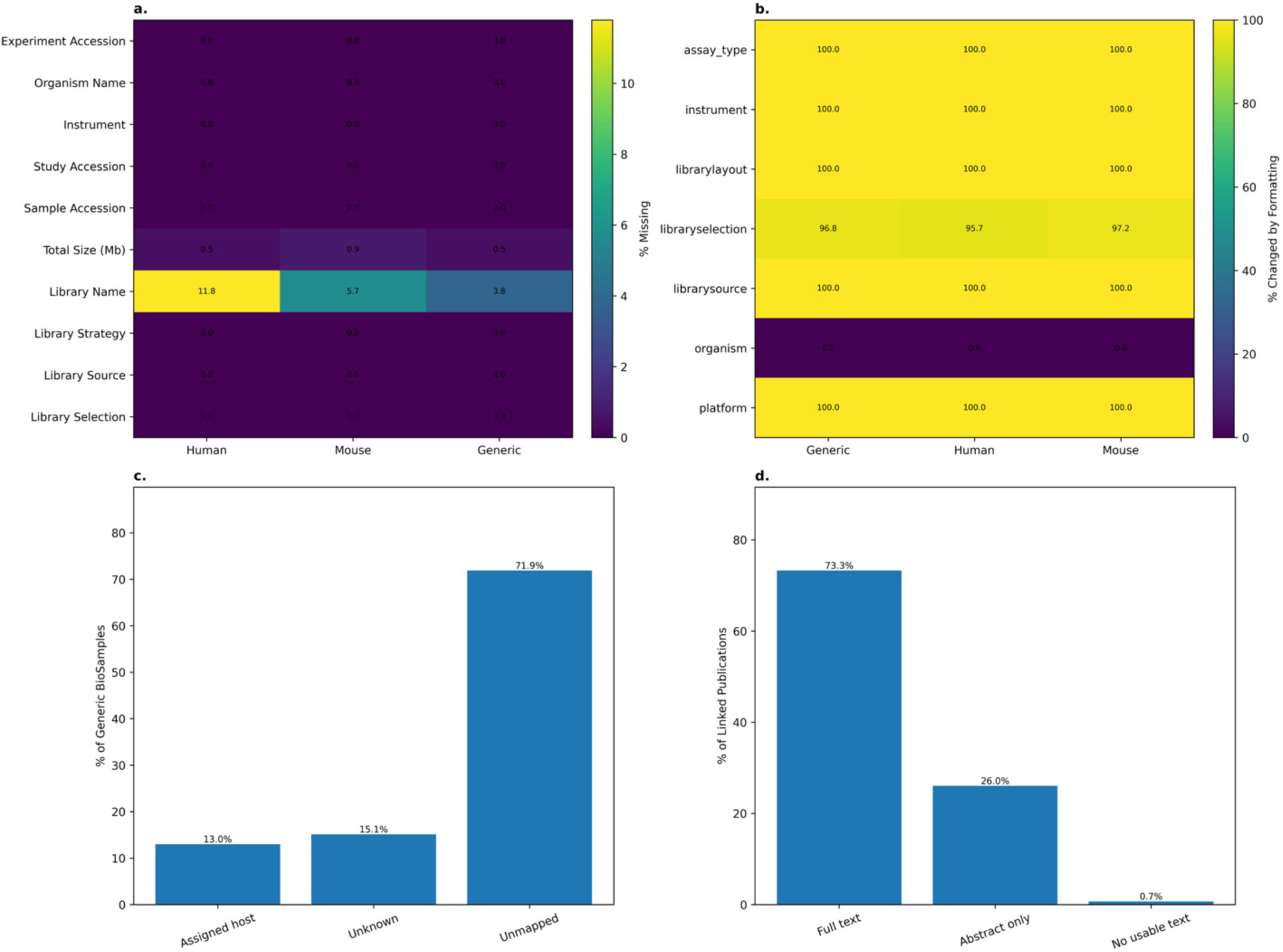
Technical metadata completeness, reporting consistency, and context recovery in public gut microbiome SRA records. (**a**) Missingness percentages for run-level metadata fields reported in **Supplemental Figure 2** across human, mouse, and generic gut metagenome cohorts. (**b**) Formatting consistency of populated metadata fields after normalization of capitalization, punctuation, spacing, and separators. (**c**) Host-attribution outcomes for generic “gut metagenome” BioSamples, showing the proportion assigned to a host, remaining unknown, or unmapped to retrievable BioSample attributes. (**d**) Publication and text-recovery outcomes for SRA-linked publications, showing the proportion with full text, abstract-only text, or no usable text available. Together, these analyses distinguish technical metadata completeness from the additional biological context and interoperability required for large-scale reuse.

However, high completeness of technical fields did not necessarily indicate that the metadata were fully standardized or biologically informative. Formatting analysis showed that most core technical categories were represented by a limited number of values, but some fields still contained substantial variation in raw terms. Instrument metadata contained 59 unique raw values in the human cohort, 65 in the mouse cohort, and 67 in the generic cohort, reflecting differences in how sequencing systems are represented across records (**Figure 5b**). Library selection and assay type also showed greater diversity than fields such as organism or library layout. These patterns suggest that even when fields are populated, additional cleaning and harmonization may be required before metadata can be used consistently across large-scale analyses.

The distinction between technical completeness and biological usefulness was especially clear in the generic “gut metagenome” cohort. Among 515,441 generic BioSamples, host identity could be assigned for only 67,005 records, corresponding to an overall host-attribution success rate of 13.00% (**Figure 5c**). BioSample attributes were retrieved for 145,000 records, and among those retrieved records, 46.21% could be assigned to a host, while 53.79% remained unknown after parsing. An additional 71.87% of generic BioSamples were unmapped to retrievable attributes in this analysis. These results show that generic gut metagenome records represent a biologically heterogeneous group and that host identity is often not recoverable from structured metadata alone.

Publication linkage provided another layer of biological context but was also incomplete at the archive level. Of 17,958 BioProjects in the analyzed dataset, 1,012 were linked to at least one PubMed identifier, corresponding to a BioProject linkage rate of 5.64%. Similarly, 545 of 18,020 SRA Studies were linked to at least one PubMed identifier, corresponding to a linkage rate of 3.02%. Among the 718 unique linked publications identified, usable text was recovered for nearly all records, including full text for 526 publications and abstract-only text for 187 publications (**Figure 5d**). Thus, once publications were linked, text recovery was high; however, only a small fraction of BioProjects and SRA Studies had direct publication links available.

Together, these results demonstrate that SRA gut microbiome records are technically well populated but not uniformly analysis ready. Basic sequencing metadata are largely complete, yet many variables needed for biological interpretation, such as host identity, phenotype, experimental design, and study outcomes, are not consistently encoded in structured run-level fields. Recovering this information often requires additional BioSample parsing, publication linkage, and full-text or abstract mining. This added layer of retrieval and harmonization complicates large-scale reuse, especially because publication links and full-text availability are not uniformly available across the archive.

### Host Attribution within Generic Gut Metagenome Records

Generic “gut metagenome” records were further analyzed to determine whether host identity could be inferred from available BioSample attribute metadata. Of the 515,441 generic BioSamples, 370,441 records (71.87%) could not be mapped to retrievable BioSample attribute metadata in this analysis, limiting host-level interpretation. Among the remaining records, 77,995 BioSamples (15.13% of the full generic cohort) had retrievable metadata but lacked sufficient information for host assignment and were classified as unknown.

Host identity could be assigned for 67,005 BioSamples, representing 13.00% of the full generic cohort. Human-associated samples were the largest identifiable host category, accounting for 47,564 BioSamples (9.23%), followed by mouse samples (10,123 BioSamples; 1.96%). Smaller identifiable host groups included dog, sheep, fish, pig, cow, rat, and chicken, each representing less than 1% of the generic cohort (**Figure 6**). These results show that records labeled broadly as “gut metagenome” represent a heterogeneous mixture of host organisms, but host identity is often unavailable, inconsistently recorded, or not recoverable from structured metadata alone. This limits automated cohort construction and demonstrates the need for additional BioSample parsing, metadata harmonization, and, in some cases, literature-based context recovery.

**Figure 6.**
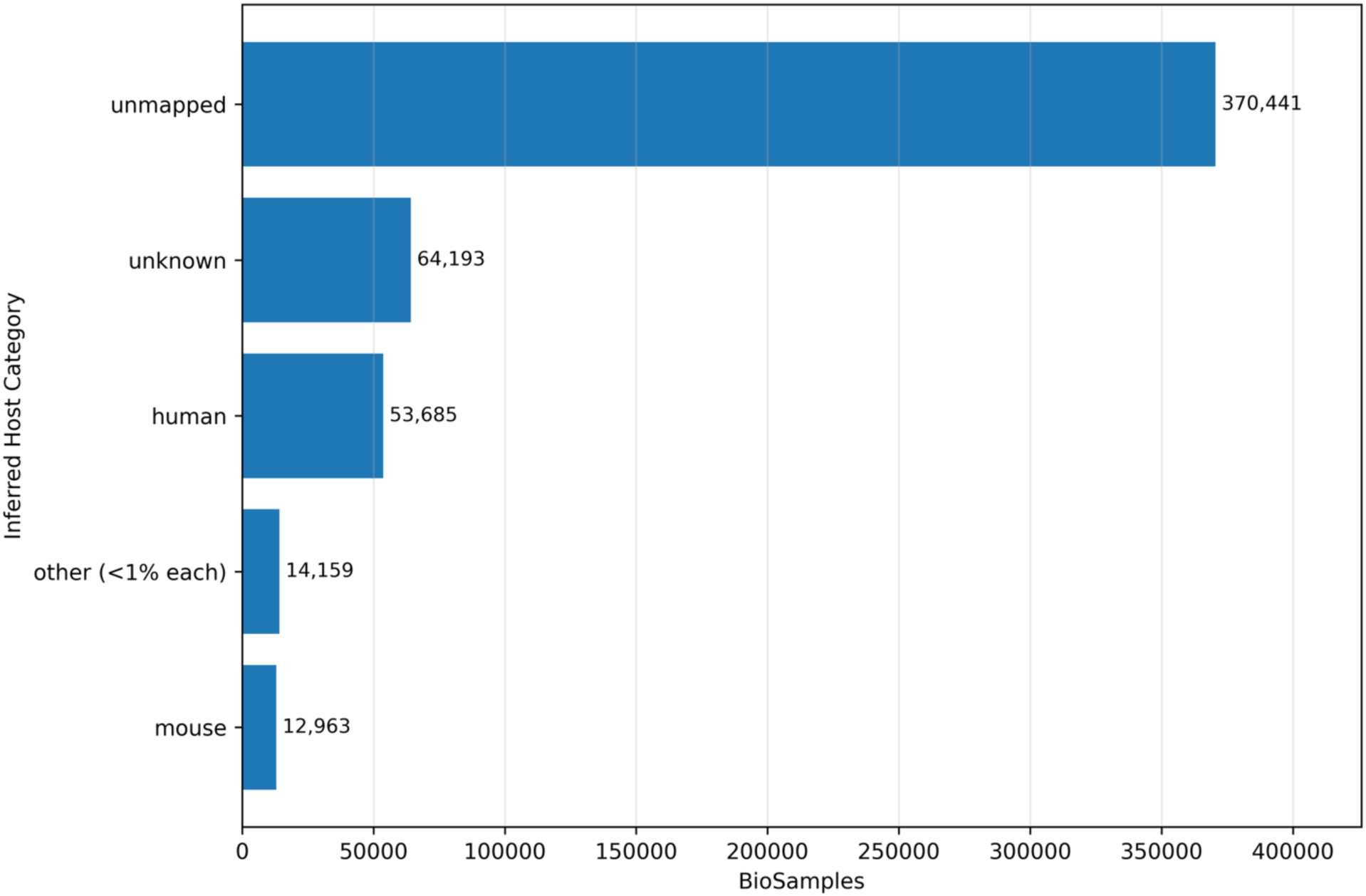
Partial Host Composition for General Gut Microbiome.

### Structural Topic Modeling of Gut Microbiome Research Literature

Topic modeling of literature linked to SRA BioProject and SRA Study accessions identified multiple temporally variable themes in gut microbiome research (**Figure 7**). The final refit structural topic model retained 20 topics, all of which are summarized with top words and generated labels in **Supplemental Figure 3**. For clarity, the main text focuses on the five most prominent and interpretable topics based on prevalence and relevance to the major research trends observed across the archive.

**Figure 7.**
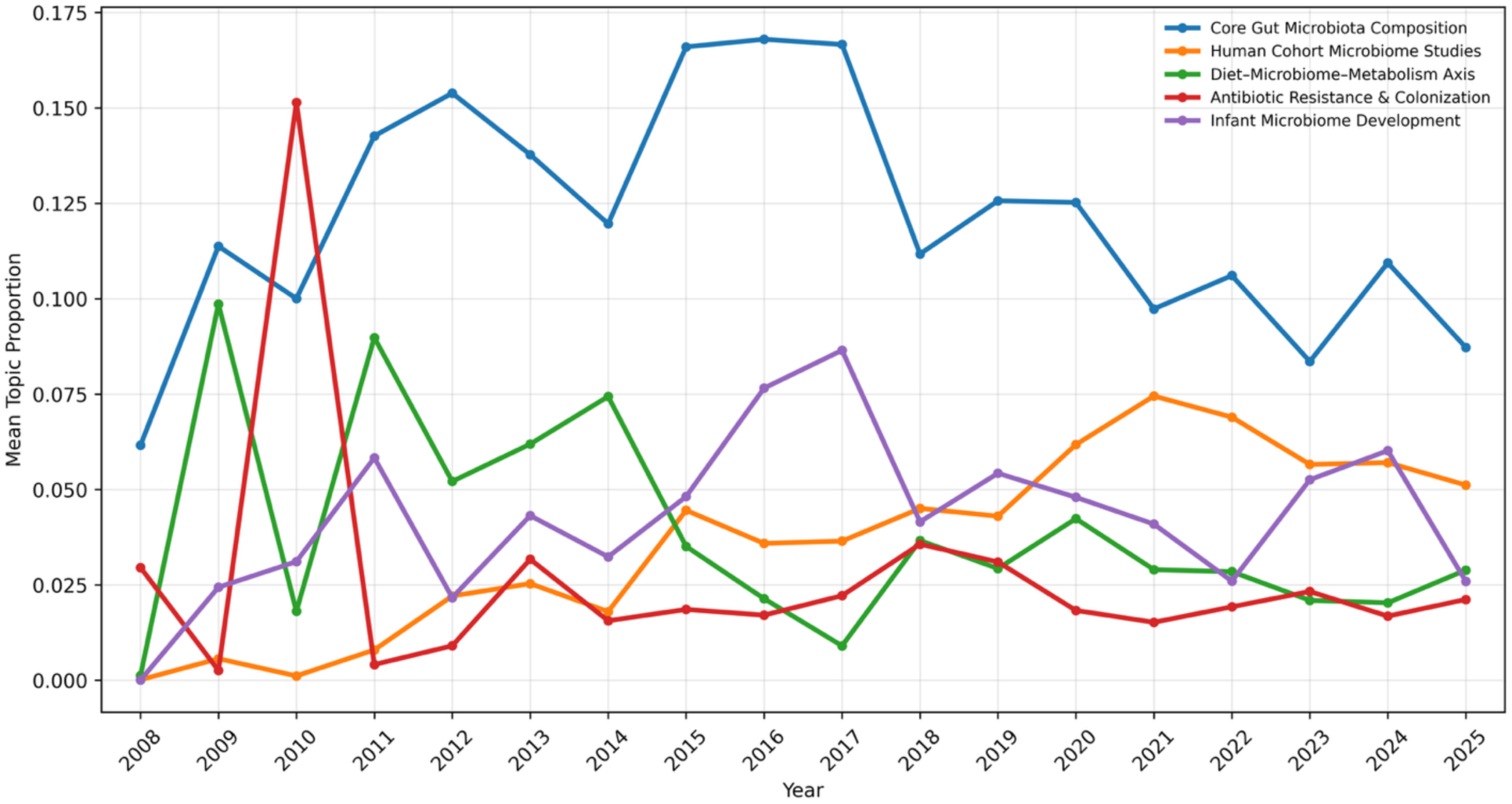
Topic prevalence was visualized from 2008 through 2025, corresponding to the main analysis period used for SRA metadata and topic-platform comparisons.

Core gut microbiota composition remained one of the most consistently represented topics across the study period. Its mean topic prevalence was 0.084 in publications from 2015 or earlier and 0.102 in publications from 2020 or later, representing a modest 1.20-fold increase; however, this temporal trend was not statistically significant (Spearman ρ = 0.31, p = 0.167). This suggests that composition-focused research remained a persistent foundation of the field rather than showing a strong directional shift.

In contrast, human cohort microbiome studies increased substantially over time. Mean prevalence for this topic increased from 0.024 in the early period to 0.062 in the recent period, corresponding to a 2.53-fold increase. This increase was supported by a strong positive association with publication year (Spearman ρ = 0.68, p < 0.001). Infant microbiome development also increased over time, rising from 0.028 in the early period to 0.042 in the recent period, with a significant positive temporal association (Spearman ρ = 0.56, p = 0.007). In contrast, diet–microbiome–metabolism and antibiotic resistance/colonization topics did not show significant monotonic increases over time. Together, these results indicate that gut microbiome research has retained a strong focus on community composition while expanding toward larger human cohort studies and developmental microbiome questions.

Topic prevalence also differed across five-year intervals. Core gut microbiota composition was highly represented across all periods, with the highest mean prevalence observed during 2015-2020. Human cohort microbiome studies showed a consistent increase across intervals, rising from 0.003 during 2005-2010 to 0.062 during 2020-2025. Infant microbiome development increased from 0.012 during 2005-2010 to 0.061 during 2015-2020 before decreasing to 0.042 during 2020-2025. These interval-based patterns suggest that the field broadened over time, with increasing representation of human population-level research and selected developmental topics alongside continued interest in core microbiome composition.

Associations between research topics and sequencing platforms were evaluated across the same time intervals to assess how thematic shifts coincided with technological change (**Figure 8**). During 2005-2010, the topic-platform associations were entirely linked to LS454 because LS454 accounted for all platform-classified runs in that interval. During 2010-2015, platform representation shifted toward Illumina, which accounted for 62.31% of runs, while LS454 still represented 37.58%. By 2015-2020, Illumina accounted for 91.19% of runs, and by 2020-2025 it increased further to 94.62%. Over the same periods, LS454 declined to 6.68% and then 0.99% of runs, respectively. Smaller contributions from Ion Torrent, PacBio SMRT, and Oxford Nanopore appeared in later intervals, reflecting gradual diversification of sequencing approaches.

**Figure 8.**
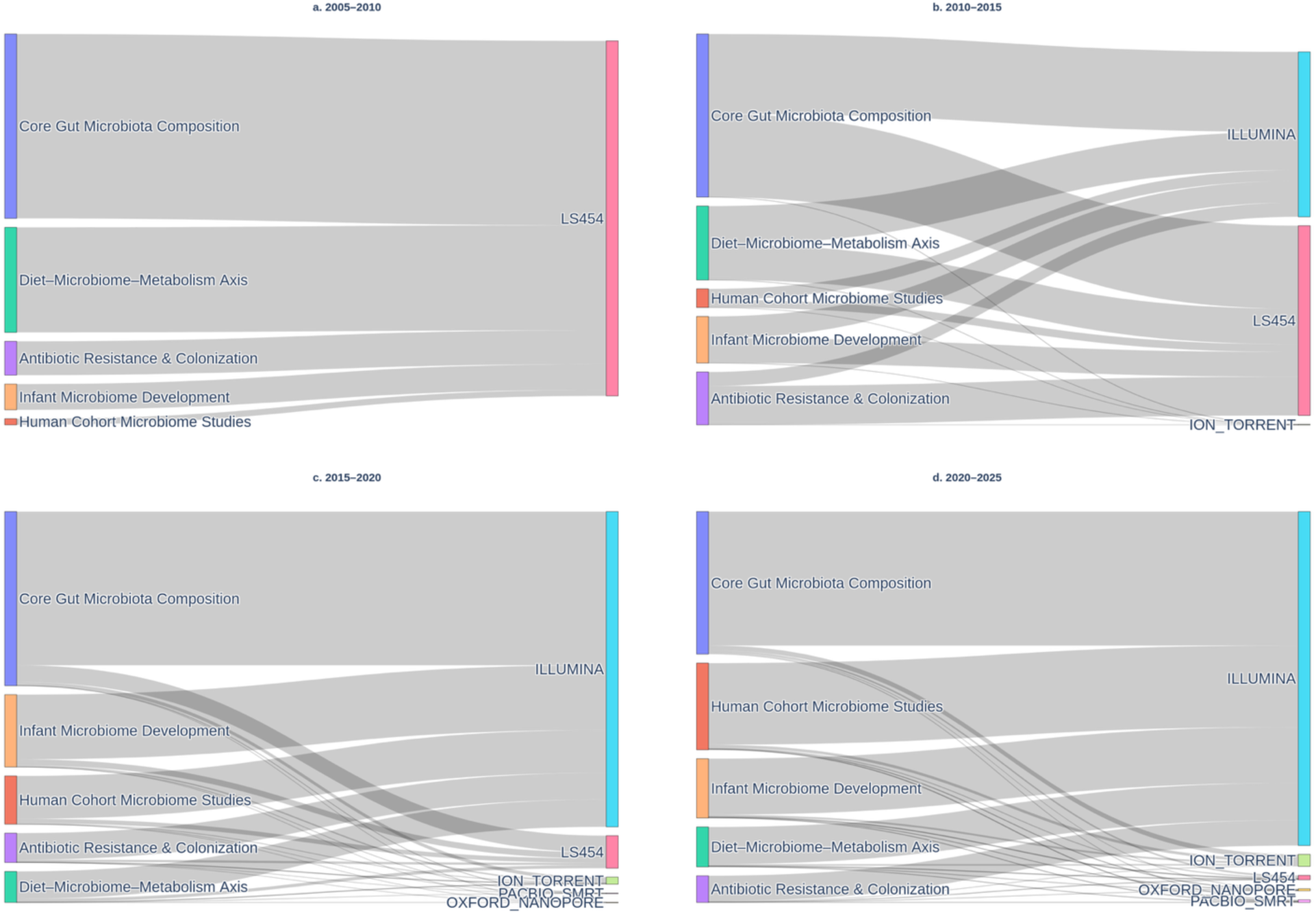
Temporal shifts in associations between gut microbiome research topics and sequencing platforms. Sankey plots depict the relationships between major gut microbiome research themes and sequencing platforms across four time intervals: **(a)** 2005-2010, **(b)** 2010-2015, **(c)** 2015-2020, and **(d)** 2020-2025. Flow widths represent the relative strength of association between topic prevalence and platform usage within each period. The figure shows an early predominance of **LS454** in foundational microbiome studies, followed by a transition toward widespread **Illumina** dominance in later periods, with smaller contributions from **Ion Torrent, PacBio SMRT,** and **Oxford Nanopore** emerging over time. These patterns illustrate how changes in sequencing technology coincided with shifts in the biological focus of gut microbiome research.

These topic-platform patterns should be interpreted as period-level associations rather than direct causal relationships between individual publications and specific sequencing platforms. The results show that shifts in research themes occurred alongside major changes in sequencing technology, especially the transition from early LS454-based studies to broad Illumina dominance. This supports the interpretation that technological adoption and evolving biological questions developed together as public gut microbiome research expanded in scale and scope.

## Discussion

### Public Gut Microbiome Archives Are Large Discovery Resources but Require Cohort-Level Harmonization

This study provides a structural characterization of publicly available gut microbiome sequencing metadata within the NCBI SRA. Rather than viewing public sequencing archives only as collections of previously published data, these resources can be understood as large-scale discovery datasets that may support cross-study comparison, model development, and hypothesis generation. Our analysis shows that gut microbiome data in the SRA have grown substantially over time, especially among human-associated datasets (**Figure 2a**). However, the same scale that makes these data valuable also creates challenges for reuse, including inconsistent metadata reporting, heterogeneous host representation, and uneven project-level sequencing concentration.

One of the clearest examples of this challenge was the heterogeneity within records labeled as “gut metagenome.” Host attribution analysis showed that these records represent a mixture of human, mouse, and other animal-associated samples (**Figure 6**). Human-associated samples made up the largest identifiable subset, accounting for 47,564 BioSamples, or 9.23% of all generic gut metagenome BioSamples. Among records that could be assigned to a host, human-associated samples represented approximately 71.0% of assigned records (**Figure 6**). However, 71.87% of BioSamples either lacked sufficient metadata for host assignment or could not be mapped to retrievable BioSample attributes, limiting fully automated reuse and comparison across studies (**Figure 6**). Similar problems have been reported in broader analyses of BioSample metadata, where inconsistent field names, uncontrolled values, and incomplete annotations reduce the ability to search, filter, and compare biological samples across repositories^5^. These results show that organism labels alone are not enough to define biologically coherent cohorts. They also point to a clear opportunity: with improved metadata harmonization and controlled sample annotations, many archived records could become more useful for integrative microbiome analysis. This interpretation is consistent with prior work showing that BioSample metadata often contain nonstandard field names and values that can impede search and secondary reuse, as well as work demonstrating that expanded metadata coverage can improve the usability of SRA-linked BioSample records^10,21^.

The metadata completeness analysis further supports this distinction between technical completeness and analysis readiness. Core technical fields were highly complete, with most evaluated variables missing in fewer than 1% of records (**Figure 5a**). However, completeness did not guarantee consistency, biological richness, or interoperability. Many biologically important variables, including host identity, phenotype, disease status, sampling context, and study outcomes, were not consistently encoded in standardized, machine-readable run-level fields. Instead, this information often remained distributed across free-text BioSample attributes or linked publications. Publication linkage and full-text availability were also incomplete, with only 5.64% of BioProjects in the dataset linked to at least one PubMed identifier, creating an additional barrier to automated recovery of biological context (**Figure 5d**). These findings suggest that the archive is technically well populated but not uniformly analysis-ready, because large-scale reuse requires not only complete sequencing descriptors but also harmonized biological context that can be compared across studies (**Figure 5**).

Our unit-of-analysis assessment further revealed that SRA data are hierarchically structured, with sequencing runs nested within BioSamples and BioProjects. Although the majority of BioSamples were represented by a single sequencing run, right-skewed distributions indicated that a small subset of BioSamples and BioProjects contributed disproportionately large numbers of sequencing runs (**Figure 2b, 2c**). This structure is consistent with broader concerns in microbiome meta-analysis, where study-level effects, batch effects, and uneven representation across studies can influence downstream biological interpretation. Prior microbiome meta-analyses have shown that study effects can confound disease-associated microbiome signatures and that batch effects must be considered when integrating microbiome datasets across cohorts^1,22,23^. In this study, project-level concentration was particularly pronounced in human-associated datasets and likely reflects the increasing role of large cohort and consortium-based studies in gut microbiome research, such as the Human Microbiome Project and the American Gut Project. The Human Microbiome Project analyzed 4,788 specimens from 242 screened adults, while the American Gut Project reported microbial sequence data from 15,096 samples from 11,336 participants, totaling more than 467 million 16S rRNA V4 reads^3^. These studies were developed to capture population-level variation in microbial communities across host factors, environments, lifestyles, and disease states, but their scale also means that a small number of projects can strongly shape the composition of public archives. This structure does not reduce the value of public microbiome data, but it does mean that downstream analyses should account for study-level clustering, sampling imbalance, and potential batch effects.

Technological evolution also played a defining role in shaping the available dataset. Sequencing depth increased substantially over time, accompanied by expanding variability across studies, reflecting increasing heterogeneity in experimental design (**Figure 2d**). Early microbiome studies often used 454 pyrosequencing because it provided relatively long reads for marker-gene sequencing, but the platform was gradually replaced as Illumina instruments became more cost-effective, higher throughput, and more accurate for large-scale sequencing studies^24,25^. Illumina platforms rapidly became dominant across all cohorts, likely because they allowed researchers to sequence larger sample sets at lower cost while relying on widely available library preparation methods and mature bioinformatics workflows (**Figure 3**). This dominance is also consistent with shotgun metagenomics workflows, where Illumina has remained the predominant platform due to its high output, accuracy, and broad availability^26^.

However, platform consistency does not eliminate all technical heterogeneity. For amplicon-based microbiome studies, primer choice and the targeted 16S rRNA variable region can strongly affect taxonomic profiles, yet these details are not always represented in standardized SRA run-level metadata. In many cases, information about the amplified region or primer set may be embedded in library names, BioSample attributes, experiment descriptions, or linked publications rather than encoded in a consistently searchable field. This further limits large-scale reuse because technically similar Illumina datasets may still differ in marker-gene region, primer design, and library preparation choices.

More recently, long-read platforms such as PacBio and Oxford Nanopore have emerged at lower frequencies, reflecting growing interest in improved assembly, strain-level resolution, and longer marker-gene reads (**Figure 3**). Together, these trends suggest a transition from early lower-throughput amplicon-focused studies toward larger, deeper, and more functionally resolved metagenomic datasets (**Figures 2d, 3, and 4**). Despite these advances, metadata completeness varied across fields, and technical completeness did not always translate into biological interpretability. This reinforces the need for systematic data harmonization approaches that preserve the value of public sequencing data while making them more comparable across studies.

### Research Themes Shift Toward Human Cohorts While Remaining Constrained by Data Structure and Technology

Structural topic modeling showed that technological and structural changes in public gut microbiome data were accompanied by shifts in the thematic focus of the field. Core gut microbiota composition remained consistently represented across the study period, suggesting that community profiling continues to serve as a foundation for gut microbiome research (**Figure 7**). In contrast, human cohort microbiome studies increased substantially in recent years, rising from a mean topic prevalence of 0.024 in publications from 2015 or earlier to 0.062 in publications from 2020 or later. This trend was statistically supported by a positive association between topic prevalence and publication year (**Figure 7**). Infant microbiome development also increased over time, indicating growing interest in early-life microbial assembly and host development (**Figure 7**). The full set of retained topics from the final topic model is provided in **Supplemental Figure 3**.

Diet-microbiome-metabolism themes were also represented in the topic model, but they did not show a significant increase over time in this SRA-linked literature subset (**Figure 7**). This distinction is important. The broader microbiome literature clearly supports diet and metabolism as major areas of biological and translational importance, but our topic model suggests that, within publications linked to the analyzed SRA records, diet-and-metabolism-related themes were present rather than expanding monotonically. Prior studies have shown that diet can rapidly alter gut microbial community structure and function, and broader reviews and systematic reviews have connected gut microbial variation to obesity, insulin resistance, type 2 diabetes, metabolic syndrome, and other metabolic outcomes^27–30^. Therefore, the topic-modeling results should be interpreted as describing the structure of SRA-linked literature rather than the full scope of diet-microbiome research.

These thematic shifts are especially relevant for AI-driven microbiome research because foundation models depend on data that are not only large, but also structured, harmonized, and biologically interpretable. The FAIR principles provide an important starting point by emphasizing that data should be findable, accessible, interoperable, and reusable^31^. However, FAIR data are not automatically AI-ready. For microbiome data to support reliable large-scale modeling and automated biological reasoning, metadata must also be sufficiently standardized, machine-readable, and connected to biological context. In the present analysis, many run-level technical fields were highly complete, but biologically important information often remained distributed across free-text BioSample attributes and linked publications (**Figure 5**). This suggests that public gut microbiome datasets are moving toward FAIR reuse but still require additional harmonization before they can fully support AI-ready workflows.

Further analysis showed that research themes were associated with sequencing technology. Illumina platforms were broadly associated with all major topics, reflecting their dominant role in microbiome sequencing, while earlier LS454-based studies were concentrated in the earliest period of the archive (**Figure 8**). More recently, long-read platforms such as PacBio and Oxford Nanopore appeared at lower frequencies (**Figure 8**). These platforms are increasingly useful because improvements in read accuracy, throughput, and bioinformatic workflows can support more contiguous assemblies, metagenome-assembled genome recovery, strain-level analysis, and improved functional interpretation^32–34^. In this context, the modest association between long-read platforms and functionally oriented topics may reflect growing interest in moving beyond community composition toward microbial function, genome reconstruction, and host–microbe mechanisms. However, these patterns should be interpreted as period-level associations rather than direct causal links between individual sequencing platforms and research questions.

Host system was also associated with research focus. Human-associated datasets were more closely aligned with population-level and cohort-based themes, likely because human microbiome studies are often designed to identify associations between microbial variation and diet, disease status, medication exposure, demographic factors, or metabolic health. Mouse-associated datasets, in contrast, are often used for mechanistic and experimental studies because animal models allow controlled perturbations, longitudinal sampling, gnotobiotic experiments, and causal testing that are not usually feasible in human cohorts^35–38^. This distinction is important for future data reuse because human and mouse datasets provide different types of biological evidence. Human datasets can support population-level modeling and clinical prediction, while mouse datasets can help test mechanisms, intervention responses, and host–microbe causality.

Recognizing this structure can help researchers build more intentional training datasets for microbiome foundation models. Rather than combining all gut microbiome records as if they represent the same type of evidence, future models could stratify or label training examples by host system, study design, sequencing platform, and biological context. Human cohort datasets could be used to learn broad population-level patterns and phenotype associations, while mouse and other experimental datasets could provide examples of controlled perturbation, intervention response, and mechanistic inference. Separating these evidence types during model development could reduce confounding, improve interpretability, and allow downstream models or agentic systems to distinguish association-rich observations from experimentally supported mechanisms.

### Public Gut Microbiome Data are AI-Relevant but Require Harmonization Before Foundation Model Use

Collectively, these findings highlight a central paradox: publicly archived microbiome sequencing data represent an unprecedented resource for discovery, but structural limitations in metadata reporting, dataset organization, and study representation constrain their immediate reuse. These limitations should not be viewed only as barriers. Instead, they identify specific points where curation, harmonization, and AI-based approaches can improve the value of data that already exist. Addressing this gap will require analytical frameworks that can integrate heterogeneous datasets, resolve inconsistencies, and extract meaningful biological signals across studies.

Foundation models represent a promising approach to this challenge because they are designed to learn generalizable representations from large, heterogeneous datasets and can be adapted to multiple downstream tasks^39,40^. Similar strategies are already emerging in microbiome and metagenomics research. Curated resources such as curatedMetagenomicData and MicrobiomeHD have demonstrated the value of reprocessing public microbiome datasets through standardized pipelines and linking them to harmonized participant or disease metadata^2,41^. These resources have enabled cross-study microbiome meta-analyses and machine-learning studies, but they also illustrate the amount of manual curation needed before public datasets can be reused reliably. More recent work using deep learning, transfer learning, and metagenomic foundation models further supports the idea that large-scale microbial sequence and abundance data can be used to learn reusable biological representations^13,42,43^. However, many of these approaches depend on curated inputs, standardized features, or selected datasets, meaning that the upstream problem of archive-scale metadata harmonization remains unresolved.

A major takeaway from this study is that the NCBI SRA already contains many of the components needed to support microbiome foundation model development, including extensive sequencing data, diverse host systems, linked literature, and large numbers of records generated using relatively uniform sequencing technology. In particular, the dominance of Illumina sequencing across cohorts suggests that much of the available sequence data may be more technically comparable than expected, even when metadata annotations remain inconsistent (**Figure 3**). This creates an important opportunity: the sequence data are already present at scale, and targeted metadata harmonization could make them substantially more useful for downstream modeling. However, this also defines a key limitation of the present archive. The data are technically rich, but they are not uniformly AI-ready because many biologically important variables remain distributed across free-text BioSample attributes, inconsistent metadata fields, and linked publications rather than standardized, machine-readable fields (**Figure 5**).

This limitation also points to a practical path forward. Expanded query strategies, controlled vocabularies, and improved BioSample-level annotation could reduce the number of relevant records missed by keyword-based filtering or hidden under broad labels such as “gut metagenome.” Ontology mapping and natural language processing could improve host attribution by linking free-text terms, scientific names, taxonomic identifiers, and sample descriptors to standardized host categories. Similarly, improved dataset-to-publication linkage and full-text mining could help recover biological context that is not encoded directly in SRA run-level metadata, including disease status, phenotype, diet, medication exposure, experimental design, and study outcomes. These steps are consistent with broader recommendations that microbiome metadata must become more standardized, FAIR, and machine-actionable before they can fully support machine learning, automation, and secondary analysis^31^.

Beyond metadata harmonization, foundation models could help organize the broader landscape of host-microbe interaction data for discovery. Integrating sequencing metadata, host phenotypes, and literature-derived knowledge could support models that identify patterns across studies, prioritize comparable datasets, and generate hypotheses about host–microbe relationships. For example, a model trained on harmonized microbiome datasets could learn that related variables such as “BMI,” “body mass index,” and BMI category represent overlapping information about host metabolic status. However, this type of reasoning depends on curated training examples that connect different field names, units, and biological meanings to shared concepts. In this context, the heterogeneity observed in SRA metadata is not only a limitation. If properly curated and labeled, this diversity can become an advantage by exposing models to variation across hosts, study designs, sequencing platforms, and biological contexts.

The limitations of this study therefore align directly with future opportunities. Metadata extraction relied on keyword-based filtering and public SRA annotations, which may have missed relevant datasets or included broadly labeled records. Host inference was limited by incomplete or inconsistent BioSample attributes, but rule-based parsing still showed that a subset of generic records could be assigned to identifiable host categories (**Figure 6**). Structural topic modeling was limited to literature linked through available PubMed identifiers, which likely represents only a subset of all relevant studies. Finally, because the analyses were observational, they do not establish causal relationships between sequencing technology, dataset composition, and research trends. Even with these limitations, the results provide a useful map of where public microbiome data are concentrated and where harmonization efforts would have the greatest impact.

The central conclusion is that NCBI gut microbiome data can be used for discovery, but they require careful structuring before they can be fully leveraged for large-scale modeling. The archive is not empty, unusable, or too fragmented to be useful. Instead, it contains a large, technically rich, and increasingly uniform sequencing resource, especially through Illumina-generated data, with metadata gaps that can be targeted through systematic curation. NCBI already has the infrastructure and archived data needed to support this next stage. With improved metadata harmonization, dataset linkage, and AI-ready data representation, public gut microbiome repositories can become a foundation for microbiome-specific models that support integrative analysis, hypothesis generation, and mechanistic discovery.

