## Supplemental Figures for "From Public Archive to Reusable Resource: Characterizing Gut Microbiome Metadata in the NCBI SRA"

NCBI SRA Metadata Metrics

| Metadata Fields |
|---|
| Experiment Accession |
| Experiment Title |
| Organism Name |
| Instrument |
| Submitter |
| Study Accession |
| Study Title |
| Sample Accession |
| Sample Title |
| Total Size, Mb |
| Total RUNs |
| Total Spots |
| Total Bases |
| Library Name |
| Library Strategy |
| Library Source |
| Library Selection |

**Supplemental Figure 1.** Mandatory reporting metrics in NCBI SRA

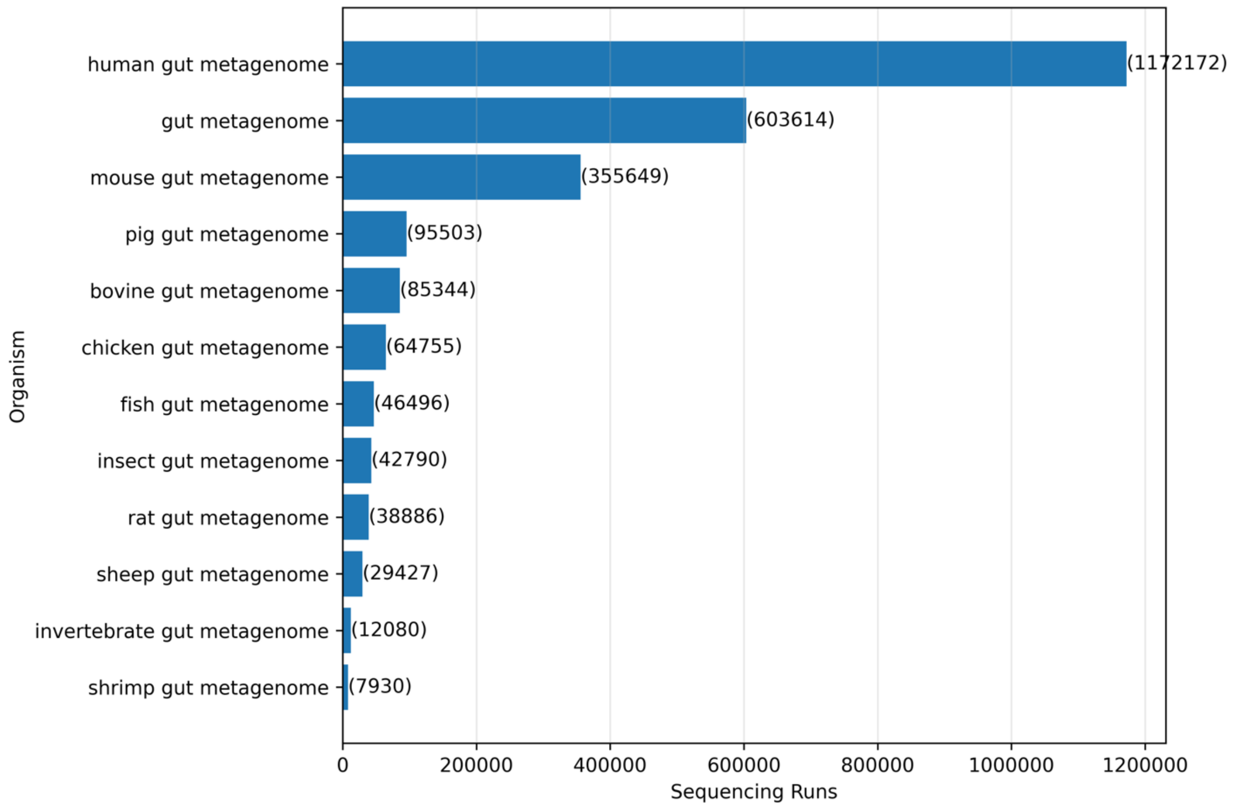


**Supplemental Figure 2.** Top Organisms in Gut Microbiome SRA

| index | topic | top_words | generated_topic_label |
|---|---|---|---|
| **0** | 0 | mice, extended, cells, after, per, pbs, tissue, neurons, days, representative, then, cre, spf, controls, lung | Mouse Immunology & Tissue Response |
| **1** | 1 | min, acid, activity, then, bacterial, cells, performed, used, 100, protein, supplementary, added, acids, metabolites, concentration | Microbial Metabolism & Biochemical Activity |
| **2** | 2 | infants, samples, age, microbiome, infant, time, between, children, abundance, gut, during, antibiotic, relative, months, microbiota | Infant Microbiome Development |
| **3** | 3 | wild, species, captive, macaques, host, laboratory, infection, identified, hosts, cats, sequences, acid, animals, humans, animal | Comparative & Evolutionary Microbiomes |
| **4** | 4 | resistance, iga, colonization, strains, strain, coli, isolates, genes, bacteria, infection, pathogen, amr, isolate, faecalis, extended | Antibiotic Resistance & Colonization |
| **5** | 5 | genes, soil, host, expression, plga, art, pregnancy, frogs, species, cpt11, frog, adult, omm12, dna, higfreo | Host Gene Expression & Molecular Interactions |
| **6** | 6 | microbiota, bacterial, between, gut, abundance, diversity, samples, microbial, otus, composition, bacteria, groups, relative, taxa, group | Core Gut Microbiota Composition |
| **7** | 7 | diet, genes, rumen, microbial, metabolism, community, animals, enzymes, gene, microbiome, dietary, their, sequences, animal, carbohydrate | Diet–Microbiome–Metabolism Axis |
| **8** | 8 | participants, between, gut, subjects, microbiota, dietary, associated, intake, bmi, individuals, variables, abundance, ibs, cohort, not | Human Cohort Microbiome Studies |
| **9** | 9 | exercise, muscle, male, weeks, female, weight, bone, microbiome, not, during, changes, intervention, alcohol, after, effects | Exercise, Physiology & Metabolic Health |
| **10** | 10 | cells, cell, tumor, cd4, cd8, tumors, responses, fetal, not, expression, immune, day, dcs, treated, cancer | Cancer Immunology & Tumor Microenvironment |
| **11** | 11 | viral, viruses, virus, phage, fungal, bacterial, virome, contigs, tmao, samples, phages, vaginal, carnitine, iav, rna | Gut Virome, Phage, and Fungal Communities |
| **12** | 12 | mice, microbiota, expression, intestinal, mouse, animals, cells, treatment, inflammation, gene, immune, not, cell, significantly, gut | Mouse Models & Intestinal Host Response |
| **13** | 13 | gut, microbiome, which, also, microbial, our, their, more, human, can, may, all, other, both, been | General Microbiome Concepts |
| **14** | 14 | 001, 0001, 005, 002, 003, 004, 006, 000, 007, 008, 009, 010, high, yes, 011 | Statistical Significance / Results Reporting |
| **15** | 15 | samples, each, sequencing, dna, reads, sample, used, all, sequences, gene, sequence, number, database, 16s, rrna | Sequencing & 16S rRNA Methods |
| **16** | 16 | diet, metabolic, levels, acid, liver, metabolism, increased, glucose, weight, dietary, fat, group, microbiota, hfd, obesity | Diet, Liver, and Metabolic Function |
| **17** | 17 | patients, disease, healthy, controls, clinical, cancer, patient, between, associated, our, ibd, microbiome, species, response, therapy | Clinical & Disease-Associated Microbiome |
| **18** | 18 | genomes, genome, supplementary, species, genes, assembly, contigs, coverage, metagenomic, mags, proteins, strain, reads, assembled, strains | Genomics & Genome-Resolved Metagenomics |
| **19** | 19 | treatment, fmt, after, donor, antibiotic, day, time, antibiotics, bla, supplementary, samples, baseline, stool, difficile, microbiota | FMT, Antibiotic Treatment & Donor Response |

**Supplemental Figure 3.** Structural Topic Model Topics and Generated Labels for SRA-Linked Gut Microbiome Literature
